# Rapid and repeatable genome evolution across three hybrid ant populations

**DOI:** 10.1101/2022.01.16.476493

**Authors:** Pierre Nouhaud, Simon H. Martin, Beatriz Portinha, Vitor C. Sousa, Jonna Kulmuni

## Abstract

Hybridization is frequent in the wild but it is unclear whether admixture events lead to predictable outcomes and if so, at what timescale. We show that selection led to correlated sorting of genetic variation in less than 50 generations in three hybrid *Formica aquilonia* × *F. polyctena* ant populations. Removal of ancestry from the species with the lowest effective population size happened repeatedly in all populations, consistent with purging of deleterious load. This process was modulated by recombination rate variation and the density of functional sites. Moreover, haplotypes with signatures of positive selection in either species were more likely to fix in hybrids. These mechanisms led to mosaic genomes with comparable ancestry proportions. Our work demonstrates predictable evolution over short timescales after admixture in nature.

---

Hybridization is widespread and has shaped the genomes of many extant species, representing a major source of evolutionary novelties *(1*–*3*). Understanding the evolution of hybrid genomes is important because it can shed light on how species barriers become established, and also on the fitness costs (e.g. incompatibilities) and benefits (e.g. heterosis) of hybridization. After admixture, local ancestry patterns within hybrid genomes are shaped by several factors (*4*). Selection may lead to the fixation (adaptive introgression, *1*) or purging of one ancestry component (incompatibilities; genetic load in one hybridizing species, *5*). Recombination rate variation can modulate the effects of selection, for instance by enabling faster purging of deleterious alleles in low-recombining regions (*5*). Admixture landscapes are also impacted by past demography and stochastic events, such as bottlenecks or initial admixture proportions. These mechanisms can lead to the fixation or near-fixation of one ancestry component at a given locus within a hybrid population, a process we refer to as *sorting* of genetic variation. Yet, few studies have investigated the interplay of different neutral and selective factors, whether they lead to repeatable sorting in hybrid genomes and at which timescale.

Here, we took advantage of multiple hybrid populations between the two wood ant species *Formica aquilonia* and *F. polyctena* to measure how rapid and predictable the evolution of admixed genomes is in the wild and identify the key factors that determine this predictability. These two species are polygynous, with up to several hundreds of queens per nest. A population is a supercolony, with dozens of interconnected nests and low relatedness between individuals (*6*). Several hybrid populations have been previously characterized in Southern Finland, and partial mitochondrial data suggests that they may have independent origins, creating an ideal test case for the outcomes of hybrid genome evolution (*7*).

We generated whole-genome sequence data from three hybrid populations (Fig. 1A, *n* = 39) and used genomes from both species collected within and outside their overlapping range (*n* = 10 per species, *8*, mean coverage: 20.6×, Table S1). Using ca. 1.6 million SNPs genome-wide, we confirmed that the hybrid populations were genetically intermediate between *F. aquilonia* and *F. polyctena* (Fig. 1, B and C, Fig. S1). Both Bunkkeri and Pikkala individuals carried distinct *F. aquilonia*-like mitotypes, while *F. polyctena*-like mitotypes were observed in the Långholmen population (Fig. 1D), where two hybrid lineages termed W and R coexist (Fig. S1, *10, 11*). Highly differentiated mitotypes between populations (Fig. 1D) and low diversity of mitotypes within a population (≤2) support a model where each hybrid population formed independently from a small number of founders.

**Figure 1.**
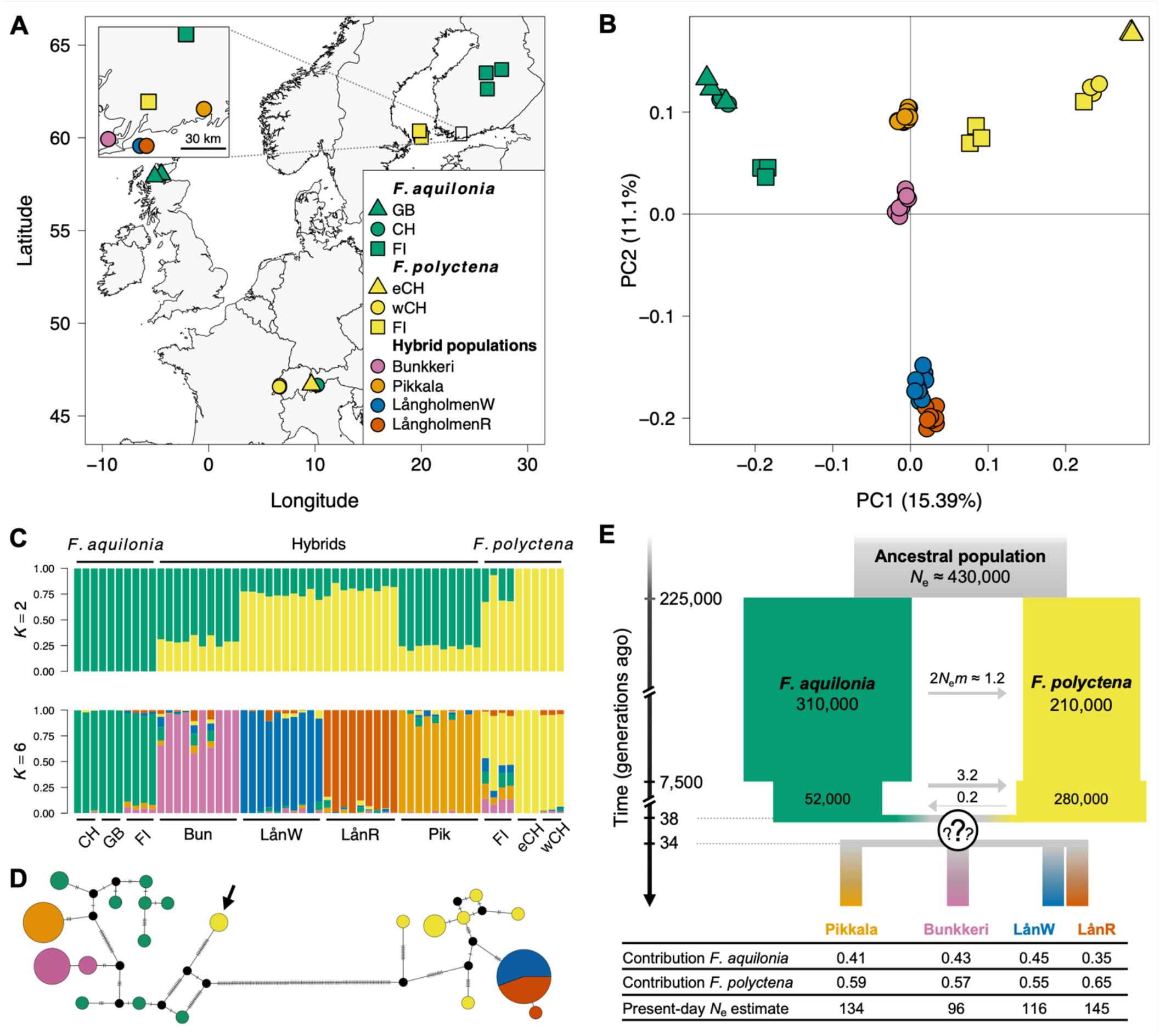
Young, independently evolving hybrid wood ant populations between *F. aquilonia* and *F. polyctena* in Southern Finland. **(A)** Sampling sites across Europe (eCH: East Switzerland, wCH: West Switzerland). **(B)** Principal component analysis of 46,886 SNPs (5 kbp-thinned, MAC ≥ 2). PC1 discriminates between both species and PC2 between hybrid populations. Colors and symbols as in panel A. **(C)** sNMF estimation of individual ancestry coefficients for *K* = 2 and *K* = 6 populations (cross-entropy criterion gives *K* = 6 as the best *K*). **(D)** Haplotype network derived from 199 mitochondrial SNPs. Circles represent haplotypes, with sizes proportional to their frequencies. Dashes indicate the number of mutational steps. The black arrow indicates two Finnish *F. polyctena* individuals carrying *F. aquilonia*-like mitotypes. **(E)** Admixture history between *F. aquilonia* and *F. polyctena* inferred through an SFS-based approach. The question marks represent the uncertainty associated with the best admixture model: both single and multiple origins scenarios show similar likelihoods, but best single origin models support separation of hybrid populations after brief periods of shared ancestry. *N*_e_: average effective population size in number of haploids, *m*: migration rate.

To elucidate the ancestry of the hybrid populations and date admixture events, we reconstructed their demographic histories using a coalescent approach based on the site-frequency spectrum [FASTSIMCOAL2, (*12*)]. Coalescent analyses support balanced admixture proportions between species (i.e., no apparent minor ancestry) and together with field observations suggest recent admixture (< 50 generations, Fig. 1E, Tables S2-S11). However, the likelihood of models with shared and independent origins were similar (Tables S7-S11), even if distinct mitochondrial haplotypes provide support for the latter (Fig. 1D). Still, parameter estimates for models that assume a shared origin indicate that the hybrid populations mostly evolved independently (on average 9.5 generations of shared ancestry since admixture, Tables S7-S11). Following the admixture event(s) no significant gene flow was inferred either between hybrid populations or between hybrids and both species (Tables S7-S11).

To investigate if evolution has resulted in repeatable admixture patterns, we mapped ancestry components along the genome independently for each hybrid population. To do this, we inferred local ancestry at 1.5 million phased SNPs using LOTER (Fig. 2A, *13*) and quantified tree topology weights in 100-SNP windows with TWISST (Fig. 2B, *14*). Hybrid populations have strongly correlated admixture landscapes along the genome (i.e., local ancestry in one population predicts the local ancestry in another population, Fig. 2C, Fig. S2, Spearman’s rank correlation tests, *P* < 10^−15^ for all population pairs). To test whether such predictability would be expected under neutrality, we used MSPRIME (*15*) to simulate neutral admixture scenarios following the best history and demographic parameters inferred for each population pair (Table S3). In all comparisons, neutral simulations led to balanced contributions of both ancestry components along the genome but did not capture the clear deviations towards either ancestry component observed locally in the genome (i.e., sorting) with our empirical data (Fig. 2D-E, two-sample Kolmogorov-Smirnov tests, *P* < 10^−15^ for all populations). Thus, correlated sorting of genetic variation among hybrid populations cannot be explained solely by the admixture scenario or their demographic history. Moreover, both SFS-based demographic modeling (Table S3) and the lack of mitochondrial haplotype sharing between hybrid populations (Fig. 1D) ruled out gene flow as a potential source for the parallelism observed. As such, other mechanisms must be invoked to explain the rapid evolution of sorted and correlated admixture landscapes in the different hybrid populations.

**Figure 2.**
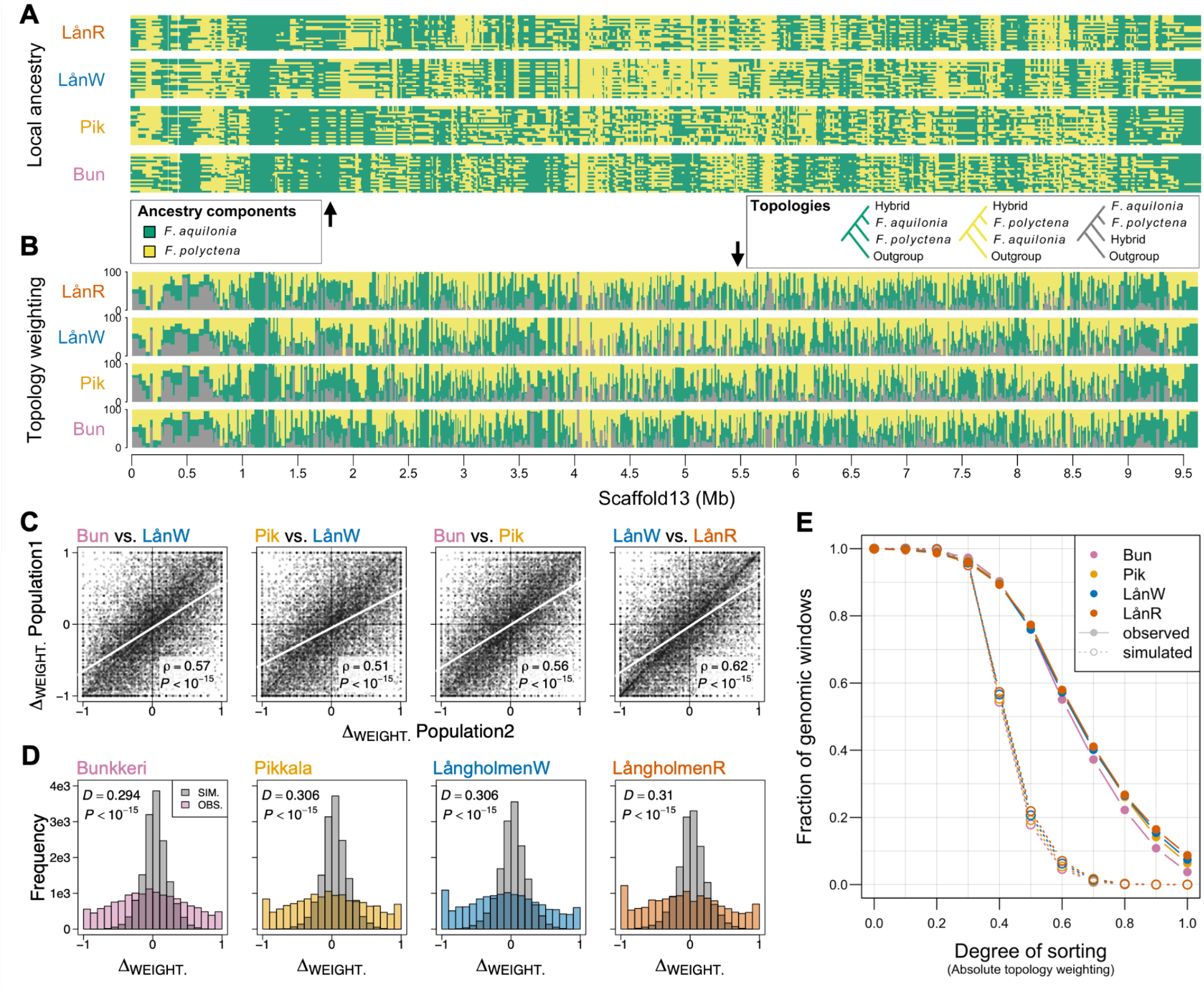
Sorting of genetic variation is more correlated than expected under neutrality across three hybrid wood ant populations. Example of **(A)** local ancestry and **(B)** topology weighting patterns inferred independently in each population along Scaffold 13. **(C)** Genome-wide comparison of topology weighting differences between each population pair (Δ_WEIGHT._: *F. aquilonia* weighting minus *F. polyctena* weighting, computed per population over 14,890 100-SNP windows, grey circles). The regression line is indicated in white. ρ, Spearman’s correlation coefficient and *P, P*-value of the Spearman’s correlation test. **(D)** Excess of extreme topology weightings in observed compared to simulated data in all populations. Distributions compared with two-sample Kolmogorov-Smirnov tests (*D*, test statistic and *P, P*-value). **(E)** A larger fraction of the genome is sorted in observed compared to simulated data in all populations (sorting measured as the absolute *F. aquilonia* or *F. polyctena* weighting).

We hypothesized that correlated sorting of genetic variation in hybrid populations is caused by selection against deleterious alleles that have accumulated in the hybridizing species with the lowest effective population size [*N*_e_, (*16*–*18*)]. This effect is expected to be stronger in gene-dense regions (*18, 19*) but also in low-recombining regions (*5*), where tighter linkage between deleterious alleles, and between neutral and deleterious alleles, leads to removal of larger tracts of ancestry. The two *Formica* species display contrasted long-term effective population sizes, with a ∼30% lower *N*_e_ in *F. polyctena* compared to *F. aquilonia* in the last 200,000 generations (*8*, Fig. 1E).

In hybrid populations, sorting (hereafter ≥90% of one ancestry component at a given locus) was faster in low-recombining regions of the genome, as well as in gene-rich regions (Fig. 3A, Fig. S3). Moreover, in low-recombining regions, the *F. aquilonia* ancestry was preferentially fixed in all populations, despite *F. polyctena* ancestry being more prevalent genome-wide (Fig. 1E). Focusing on coding SNPs, we found a significant enrichment for *F. aquilonia* ancestry genome-wide in all populations, consistent with the hypothesis that hybrids can purge the deleterious load accumulated in *F. polyctena* due to its smaller *N*_e_ (genomic permutations, *P* < 0.002 in all populations, Fig. 3B). These results support the contributions of both recombination rate variation and genetic load in promoting sorting of ancestral variation in hybrids. They also reveal that consistent sorting of parental variation can happen in less than 50 generations in small populations (Fig. 1E).

**Figure 3.**
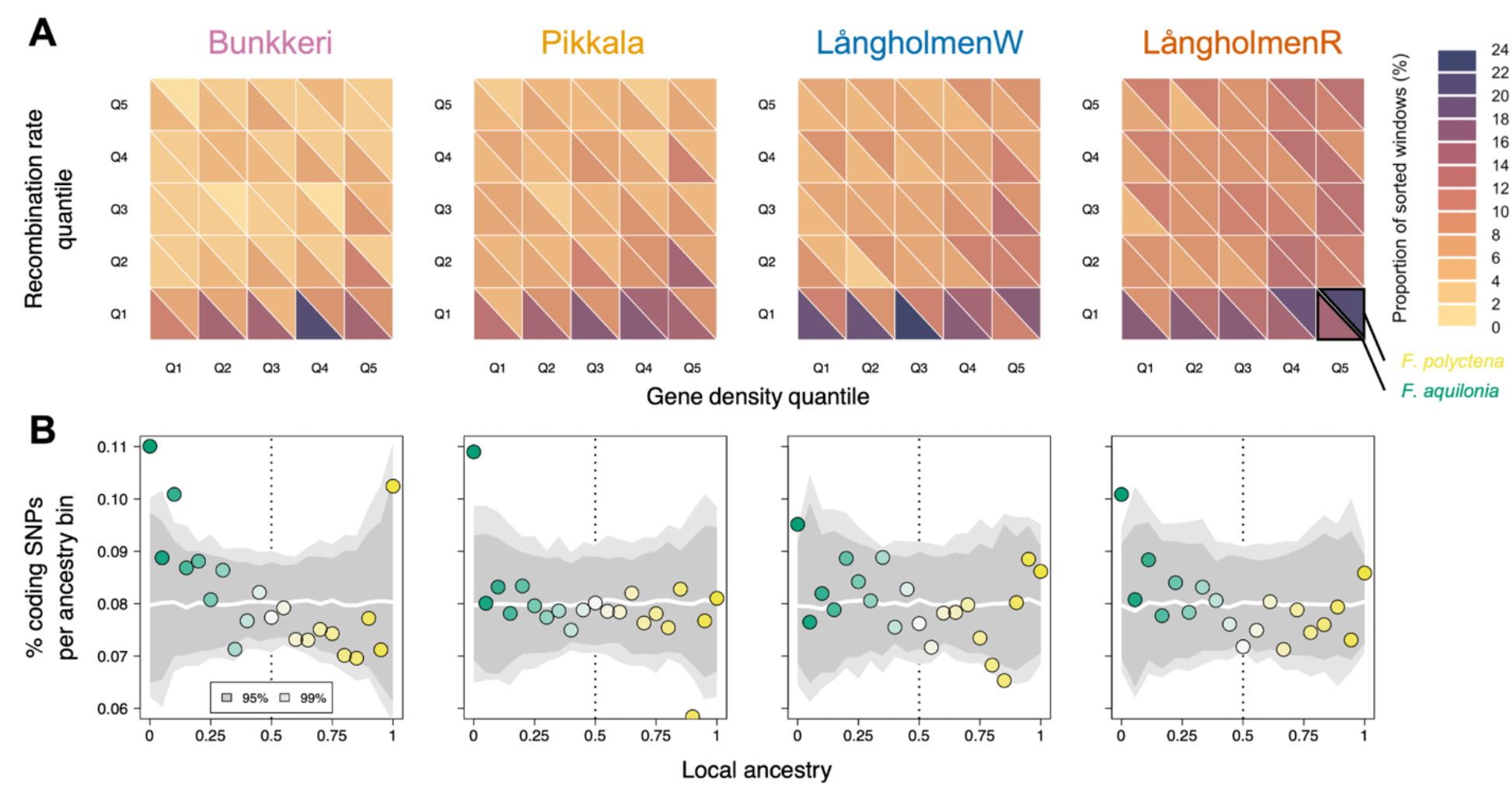
Sorting of ancestral polymorphism in hybrids is driven by recombination rate variation and genetic load. **(A)** Heatmap showing the fraction of sorted 20 kbp windows and the direction of sorting (ancestry component fixed) as a function of recombination rate and gene density quantiles in each hybrid population. **(B)** Coding regions are significantly enriched for the *F. aquilonia* ancestry component in all hybrid populations (*P* < 0.002). For each population (panel, same as A) is plotted the fraction of SNPs within CDS (y-axis) as a function of local ancestry (x-axis, 0: *F. aquilonia* allele fixed, 1: *F. polyctena* allele fixed). Confidence intervals (in grey) were obtained using 500 genomic permutations (white line: median of the permutation approach).

Positive selection could also drive correlated sorting of genetic variation across hybrid populations: advantageous alleles from either species could repeatedly sweep in distinct hybrid populations after admixture. Under this scenario, a genomic region fixed for the *F. aquilonia* ancestry component in hybrids would display signatures of selection in *F. aquilonia* individuals. To test this hypothesis, we looked for selective sweep signatures in both species with RAiSD (*21*), which quantifies changes in the SFS, levels of LD and genetic diversity along the genome through the composite sweep statistic μ (Fig. 4). Consistently sorted genomic windows in hybrids (i.e., windows fixed for either species ancestry across all hybrid populations, 1.92% of the windows overall) displayed significantly higher sweep statistics only in the species from which the ancestry component was fixed in hybrids (genomic permutations, *P* < 0.001, Fig. 4B). While recombination rate estimates were significantly lower than the rest of the genome in these consistently sorted windows (Wilcoxon test, W = 1,740,660, *P* < 10^−15^), purging of load in low recombining regions cannot explain the observation that hybrids have fixed ancestry from the species where a sweep may have occurred prior to hybridization (Fig. 4B).

**Figure 4.**
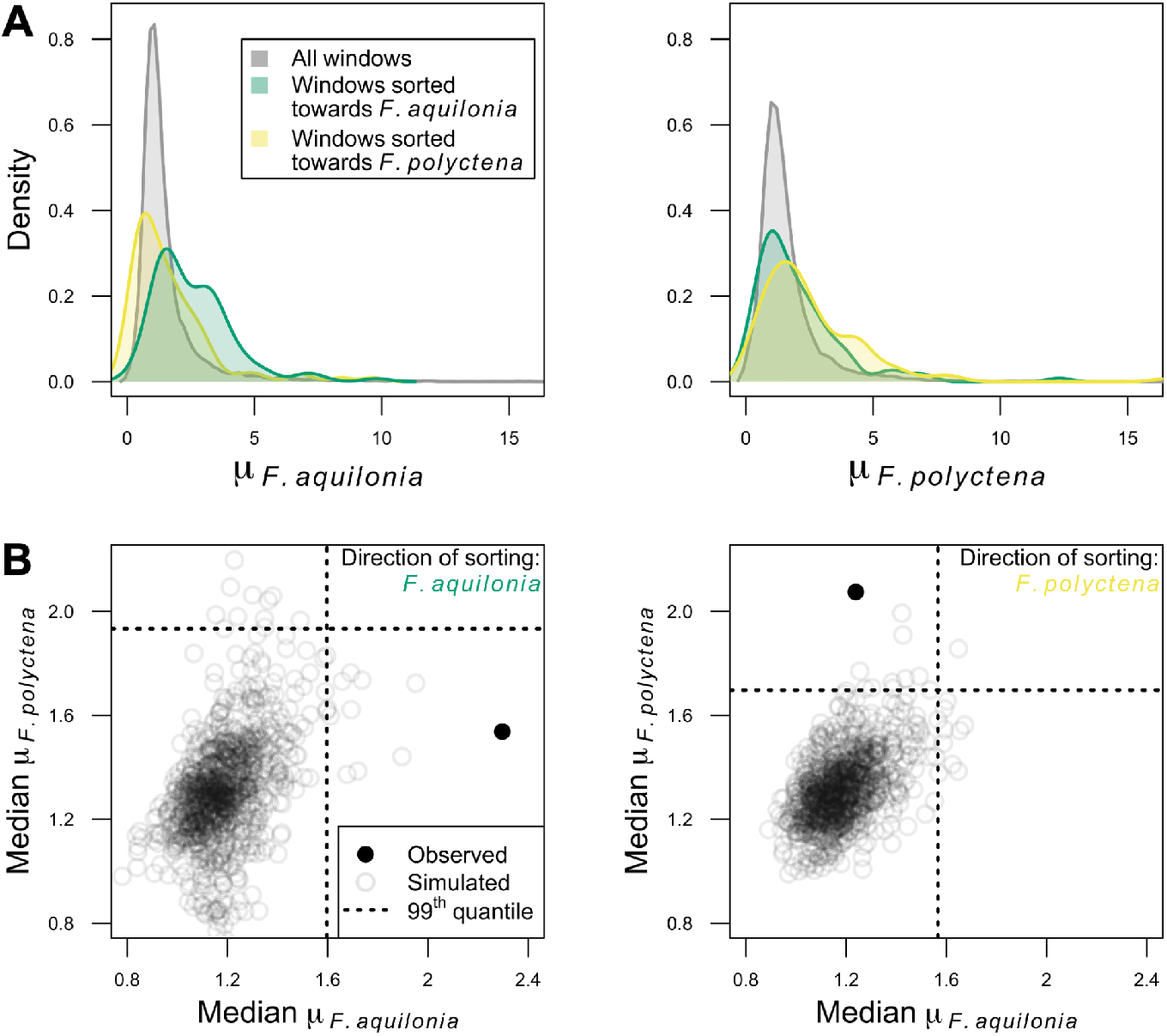
Signatures of selective sweeps in hybridizing species predict the direction of sorting in admixed genomes. **(A)** Distribution of selective sweep statistics (μ) computed over 20 kbp windows in *F. aquilonia* (left) and *F. polyctena* (right). Genome-wide μ distribution (grey) and observed values in windows fixed for either *F. aquilonia* (*n* = 104 windows, green) or *F. polyctena* ancestry components (*n* = 98, yellow) in all hybrid populations. **(B)** Windows fixed for either *F. aquilonia* (left) or *F. polyctena* (right) ancestry components in all hybrid populations are significantly enriched for high μ values in *F. aquilonia* or *F. polyctena* individuals, respectively. This suggests that a haplotype with a signature of positive selection in either species is more likely to fix in hybrids. Simulated μ values were obtained through 1,000 genomic permutations (as in *20*). Each circle represents medians computed over all consistently sorted windows (plain: observed, open: simulated).

Hybrid genomes provide powerful insights into evolution because they are exposed to strong, and often opposing, selective forces (*4*). Here we have shown that the sorting of ancestral genetic variation in hybrid genomes can occur rapidly and predictably after admixture due to both positive and purifying selection.

## MATERIAL & METHODS

### Sampling

*F. aquilonia, F. polyctena* and their Finnish hybrids are polygynous: within a nest, the reproductive effort is shared across dozens or hundreds of egg-laying queens. These two species and their Finnish hybrids are also supercolonial, with populations (i.e., supercolonies) formed by the association of several cooperating nests within a site. Polygyny and supercoloniality both result in low relatedness among individuals sampled within a given population (*22*). We sampled hybrid individuals from three populations previously mapped in Southern Finland: Långholmen (composed of two hybrid lineages R & W, *10*), Bunkkeri and Pikkala (*7*). We collected unmated queens from Bunkkeri (*n* = 10) and Långholmen (*n*_W_ = 10, *n*_R_ = 9) in Spring 2018 and workers from Pikkala (*n* = 10) in Spring 2015 (Table S1). Data generated by Portinha *et al*. (*8*) were used as *F. aquilonia* and *F. polyctena* reference panels. Briefly, it consists of ten workers (diploid females) per species sampled from several locations across Europe (Fig. 1A). *F. polyctena* samples were collected at two locations in Switzerland (East and West) and in the Åland islands (Finland), and *F. aquilonia* samples were collected in Scotland (UK), East Switzerland and Central Finland.

### DNA extraction

Both the hybrid samples generated for this study and the *F. aquilonia* and *F. polyctena* samples from Portinha *et al*. (*8*) were processed and sequenced at the same time, and all data went through the same pipeline. DNA was extracted with a Sodium Dodecyl Sulfate (SDS) protocol from whole bodies and sequencing libraries were built with NEBNext DNA Library Prep Kits (New England Biolabs) by Novogene (Hong Kong).

### DNA sequencing and read mapping

Unless stated otherwise, all software was used with default parameters. Whole-genome sequencing was carried out on Illumina Novaseq 6000 (150 base pairs, paired-end reads), targeting 15× per individual (Table S1). We trimmed raw Illumina reads and adapter sequences with TRIMMOMATIC (v0.38; parameters LEADING:3, TRAILING:3, MINLEN:36; Bolger et al., 2014) and mapped trimmed reads against the reference genome (*23*) using BWA MEM (v0.7.17, *24*). We then removed duplicates using PICARD TOOLS (v2.21.4; http://broadinstitute.github.io/picard).

### SNP calling and filtering

We called single nucleotide polymorphisms (SNPs) jointly across all samples with FREEBAYES (v1.3.1, population priors disabled with -k option, *25*) and normalized the resulting VCF file using VT (v0.5, *26*). We excluded both sites located at less than two base pairs from indels and sites supported by only Forward or Reverse reads using BCFTOOLS (v1.10, *27*). We decomposed multi-nucleotide variants using vcfallelicprimitives from VCFLIB (v1.0.1). The next steps were carried out with BCFTOOLS. Biallelic SNPs with quality equal or higher than 30 were kept. Individual genotypes with (*i*) genotype qualities lower than 30 and/or (*ii*) with depth of coverage lower than eight were coded as missing data. Sites displaying more than 50% missing data over all samples were discarded. Genotyping errors due to e.g., misaligned reads were removed using a filter based on excessive heterozygosity. To do so, we used an approach similar to Pfeifer *et al*. (*28*) and excluded sites displaying heterozygote excess after pooling all samples (P < 0.01, --hardy command from VCFTOOLS v0.1.16, *29*). We then filtered sites based on individual sequencing depth distributions, setting as missing sites where depth was lower than half or higher than twice the mean value of the individual considered. Finally, sites with more than 15% missing data over all samples were discarded. These steps led to a final data set of 1,659,532 SNPs across 59 individuals.

### Population structure

Population structure was assessed using a reduced data set of 46,896 SNPs obtained after thinning (retaining one SNP every 5 kbp) and discarding sites with minor allele count ≤ 2. We performed a Principal Component Analysis (PCA) with PLINK (v1.9, *30*) and sNMF clustering using the LEA package (v3.0.0, *31*) in R (v3.6.2, *32*). Clustering was carried for a number of ancestral components (*K*) ranging from 1 to 10, with ten iterations performed per *K* value. The lowest cross-entropy was obtained with *K* = 6 and the results of runs with the lowest cross-entropy for *K* = 2 and *K* = 6 are shown in Fig. 1C.

### Mitotype network

Mitochondrial SNPs were called separately with FREEBAYES using a frequency-based approach (--pooled-continuous option). SNP filtering was carried using the same pipeline as for the nuclear genome, which led to the identification of 199 biallelic SNPs across 59 individuals. Individual FASTA files were written using vcf-consensus and aligned with MAFFT (v7.429, *33*). The median-joining network was created using POPART (*34*).

### Demographic modeling

Before reconstructing admixture histories, we removed the 122,479 SNPs located on the third chromosome. This chromosome carries a supergene controlling whether *Formica* colonies are headed by one or multiple queens (social chromosome, *9*). Recombination reduction between the two supergene variants leads to the maintenance of ancestral polymorphisms across *Formica* species which could bias our demographic inference. The data set used for demographic modeling hence comprises 1,537,053 SNPs.

We used the composite-likelihood method implemented in FASTSIMCOAL2 (v2.6, *12*) to compare alternative demographic models to demographic parameters inferred from the site frequency spectrum (SFS) following Portinha *et al*. (*8*). We ran each model 100 times with 80 iterations per run for likelihood maximization and the expected SFS was approximated through 200,000 coalescent simulations per iteration. We assumed a mutation rate of 3.5×10^−9^ per bp per haploid genome per generation, which is an average based on estimates currently available for social insects (*35*). No population growth was allowed, but population sizes could change after admixture events and/or when migration rate changed. Generation time was assumed to be 2.5 years (*8*). Finally, we used the best speciation history inferred by Portinha *et al*. (*8*) to constrain parameter range prior to the admixture event in our demographic models. This speciation history was inferred from the same Finnish *F. aquilonia* and *F. polyctena* individuals as used in the present study. All parameter ranges are indicated in Tables S2 (three-population models) and S7 (four-population models). For each comparison, the model with the highest expected likelihood was chosen as the best model.

Field observations suggest hybrid populations may have arisen through recent admixture (ca. 50 years ago). Constraining admixture under 50 generations (ca. 125 years ago) led to models with higher expected likelihoods for all hybrid populations, and both constrained and unconstrained results are shown in Tables S3-S6.

### SFS characteristics

We built folded SFSs using minor allele frequency (MAF) and downsampled genotypes to minimize missing data. We first determined a minimum sample size across all sites (number of individuals available for resampling minus maximum number of missing data per site). We then resampled individuals in 50 kbp windows and discarded blocks where the mean distance between two consecutive SNPs within a block was < 2 bp. We estimated the number of monomorphic sites from the proportion of polymorphic sites and the total number of callable sites. The latter was obtained from each individual BAM file using MOSDEPTH (v0.2.9, *36*) and individual sequencing depth thresholds defined for SNP calling. Distinct 3D- and 4D-SFSs were built for our 3- and 4-population models, respectively to answer specific study questions (see below). In both cases, we used the single individual from each species sampled in Southern Finland as representative of their respective species. For each hybrid population, we resampled four individuals every window to build the SFSs. The 3D-SFSs contained information of both species and one focal hybrid population, while the 4D-SFSs included information of both species and two focal hybrid populations. For these latter models we analyzed four different combinations: Bunkkeri - LångholmenW, Pikkala - LångholmenW, Bunkkeri - Pikkala and finally LångholmenW - LångholmenR.

### Disentangling between secondary contact and hybridization: three-population models

For each hybrid population we tested three different scenarios which could lead to present-day admixed individuals. The first scenario is hybridization, namely an admixture event between *F. polyctena* and *F. aquilonia* where one species would contribute a genetic input of *α* into the hybrid population while the other species would contribute the remaining fraction 1 - *α*. This scenario was assessed both with and without gene flow between hybrids and either (or both) species after admixture (forward in time).

The second scenario is secondary contact, where after the speciation event hybrid ancestors would first diverge from one species, and then receive migrants from the other species. This scenario was tested in both directions [i.e., assuming both ((*F. polyctena*, hybrid), *F. aquilonia*) and ((*F. aquilonia*, hybrid), *F. polyctena*) topologies]. Gene flow was also allowed between both species before and after the split between the hybrid population and the first species.

The third and last scenario is a trifurcation model, where the two species and the hybrid ancestral population would first diverge simultaneously, after which all populations would exchange migrants at different rates. Higher migration from both species into the hybrid ancestral population would lead to admixed individuals in the current-day hybrid population.

### Disentangling between shared and independent origin of hybrid populations: four-population models

Hybrid populations that arose through a single admixture event followed by a long period of shared ancestry would have more correlated sorting of genetic variation than if they separated soon after the admixture event or arose through independent admixture events. To disentangle between these scenarios, we tested two alternative admixture models using two populations at a time. Based on the results of our three-population models, admixture times were constrained to the last 50 generations in all subsequent models. The first model is a single-origin model where *F. polyctena* contributes a proportion *α* of the genetic material of the ancestral hybrid population, with *F. aquilonia* providing the complementary 1 - *α* fraction. This ancestral hybrid population then diverges into two hybrid populations.

The second model is an independent-origins model where each hybrid population arises through a distinct admixture event, with possibly different contributions from both species (i.e., *F. polyctena* contributes a fraction *α* for the first hybrid population and *β* for the second hybrid population, and *F. aquilonia* 1 - *α* and 1 - *β*, respectively). In this second model, the order of each admixture event depends on admixture dates inferred through three-population models.

As post-admixture gene flow between hybrid populations could also lead to correlated sorting of genetic variation, we additionally tested an independent-origins-with-migration model which includes reciprocal migration between two hybrid populations after the second (most recent) admixture event.

### Population recombination rate estimation

We used iSMC (v0.0.23, *37*) to estimate population recombination rates along the genome. We hypothesized that the recombination landscape in hybrids would be an average of the recombination landscapes in both species. Using all non-Finnish individuals from both species jointly, we fitted a model of coalescence with recombination including 40 times to the most recent common ancestor (TMRCA) intervals and 10 ρ categories. Population recombination rate estimates were collected over non-overlapping 20 kbp windows, discarding windows with less than 20 SNPs.

### Haplotype estimation

Prior to mapping ancestry components, we phased all SNPs anchored on scaffolds (78.2% of the genome) with WHATSHAP (v1.0, *38*), which uses short range information contained within paired-end reads. We then performed statistical phasing and imputation with SHAPEIT (v4.1.2, *39*), using the sequencing data setting and increasing the MCMC iteration scheme as indicated in SHAPEIT documentation. The total phased data set contained 1,490,364 SNPs.

### Outgroup information

We used *F. exsecta* as an outgroup to root topologies inferred with TWISST (see below). This species belongs to a distinct species group (*F. exsecta* group) while *F. aquilonia* and *F. polyctena* both belong to the *F. rufa* group (*40*). To extract *F. exsecta* genotypes at our phased SNP loci, we mapped data previously generated by Dhaygude *et al*. (*41*) against the same reference genome used in the present study. These data consist of Illumina paired-end, 2×100 bp reads generated from a pool of 50 (haploid) males (9.89 Gbp overall, median insert size: 469 bp, ENA accession SAMN07344806). Reads were trimmed with TRIMMOMATIC using the same parameters as before, mapped with BWA MEM and deduplicated with PICARD TOOLS. We filtered proper read pairs with mapping quality ≥ 20 (86.4% of the reads, sequencing depth 14.5×) using SAMTOOLS (v1.13, *42*). We generated a pileup file (disabling per-base alignment quality computation and filtering minimum base quality ≥ 20) and finally extracted *F. exsecta* genotypes at each locus of the phased data set using a custom R script.

### Mapping ancestry components along the genome

We used three different methods to map ancestry variations along the genome, which all relied on reference panels comprising both *F. aquilonia* and *F. polyctena* individuals. As gene flow from *F. aquilonia* to *F. polyctena* in Finland prior to admixture (*8*) could bias our results, we ran all three methods after excluding the Finnish representatives of both species from our reference panels.

We performed local ancestry inference from phased data for each population independently using LOTER (v1.0, *13*), after which we averaged local ancestries over the same non-overlapping 20 kbp windows used for population recombination rate estimation.

We also recorded topology weightings in 100-SNP windows along the genome with TWISST (v0.2, *14*) using phased data. Analyzing jointly the two species, the outgroup *F. exsecta* and one hybrid population at a time, the three rooted topologies are (Fig. 2B):

1. (((*F. aquilonia*, hybrid), *F. polyctena*), outgroup),
2. (((*F. polyctena*, hybrid), *F. aquilonia*), outgroup),
3. (((*F. aquilonia, F. polyctena*), hybrid), outgroup).

Since the average weighting of the third topology was around 10% genome-wide in all hybrid populations, we measured topology weighting differences per window by subtracting the weighting of the second topology (*F. polyctena* topology) to the weighting of the first topology (*F. aquilonia* topology, Fig. 2C and D). The resulting metric, Δ_WEIGHT._, ranges from - 1 to +1 (in a given window, only the *F. polyctena* or the *F. aquilonia* topologies are inferred, respectively). When Δ_WEIGHT._ is close to zero, both topologies have similar weightings, which we interpreted as a lack of sorting of ancestral variation in hybrids.

Finally, we also performed a “naive” chromosome painting approach from non-phased genotypes. To do so, we first discarded sites with more than two missing genotypes over reference individuals, after which we identified 79,336 ancestry-informative SNPs displaying an allele frequency difference above 80% between both species.

### Coalescent simulations

We used coalescent simulations to measure sorting levels which would be expected in each hybrid population under the reconstructed admixture history. To do so, we used the parameters inferred by the best model identified for each population pair with FASTSIMCOAL2 and ran simulations in MSPRIME (v1.0.2, *15*), modeling 100 non-recombining 10 kbp blocks with a recombination rate of 10^−6^ within blocks. Each simulation was ran 100 times. Topology weightings were computed directly from tree sequences using TWISST and Δ_WEIGHT._ distributions were obtained as stated above over all 100 replicates.

### Selective sweep detection

We looked for evidence for selective sweeps independently in both hybridizing species with RAiSD (v2.9, *21*). The composite selective sweep statistics μ were estimated with default parameters using non-Finnish individuals from each species. Results were then averaged over 20 kbp non-overlapping windows in R (Fig. 4A).

### Genomic permutation approach

We tested statistical significance of both (*i*) the enrichment in *F. aquilonia* ancestry at coding SNPs (Fig. 3B) and (*ii*) the association between the direction of sorting and the evidence for selective sweeps in one species (Fig. 4B) using a similar shift-based, circular permutation scheme inspired by Yassin *et al*. (*20*).

For the first analysis, we slid the local ancestry landscape inferred by LOTER at the SNP level by 100 kbp increments (i.e., five 20 kbp windows at a time), maintaining the structure of ancestry blocks as observed in our data. For each shift replicate, we then computed the fraction of coding SNPs at each local ancestry bin genome-wide. For this analysis we ran 500 permutations and *P*-values were defined as the proportion of shift replicates in which at least a similar fraction of coding SNPs was reached as in our empirical data. The 95^th^ and 99^th^ quantiles plotted in Fig. 3B were computed over all 500 permutations.

For the second analysis, we slid the average local ancestry values computed over 20 kbp non-overlapping windows by 100 kbp increments, which also shifted the location of sorted windows. For each shift replicate, we then computed the median composite selective sweep statistic μ across all sorted windows genome-wide for each ancestry component independently. We ran 1,000 permutations and defined *P*-values as the proportion of shift replicates in which median μ values were at least as high as observed in the empirical data (Fig. 4B).

### Density of functional sites and gene content in sorted regions

The density of functional sites was computed in 20 kbp non-overlapping windows along the genome by measuring for each window the fraction of base pairs falling within coding domain sequences, which positions were extracted from the GFF annotation file (*23*). The 202 sorted windows (defined as displaying ≥90% of either species ancestry across all hybrid populations, see Fig. 4) intersected with 364 gene models, however no significant gene enrichment was detected with T_OP_GO (v1.0, *43*).

## Acknowledgments

We thank G. Barroso for assistance with iSMC, the SpecIAnt group for feedback and CSC – IT Center for Science, Finland, for computational resources. This work was performed under the Global Ant Genomic Alliance.

## Funding

HiLIFE (JK)

Academy of Finland no. 309580 (JK)

Fundação Ciência e Tecnologia CEECINST/00032/2018/CP1523/CT0008 (VCS)

Fundação Ciência e Tecnologia CEECIND/02391/2017 (VCS)

## Author contributions

Conceptualization - PN, SHM, JK.

Methodology - PN, SHM, BP, VCS, JK.

Funding acquisition - JK.

Investigation - PN, BP.

Project administration - JK.

Resources - PN, SHM, VCS, JK.

Supervision - PN, JK.

Visualization - PN.

Writing - original draft - PN, SHM, JK.

Writing - review & editing - PN, SHM, BP, VCS, JK.

All authors approved the manuscript before submission.

## Data and materials availability

Data will be made available pending acceptance of the manuscript.

## SUPPLEMENTARY MATERIALS

Tables S1 – S11

Figures S1 – S3

References (22 – 43)

