## Supplementary Figures for "Rapid and repeatable genome evolution across three hybrid ant populations"

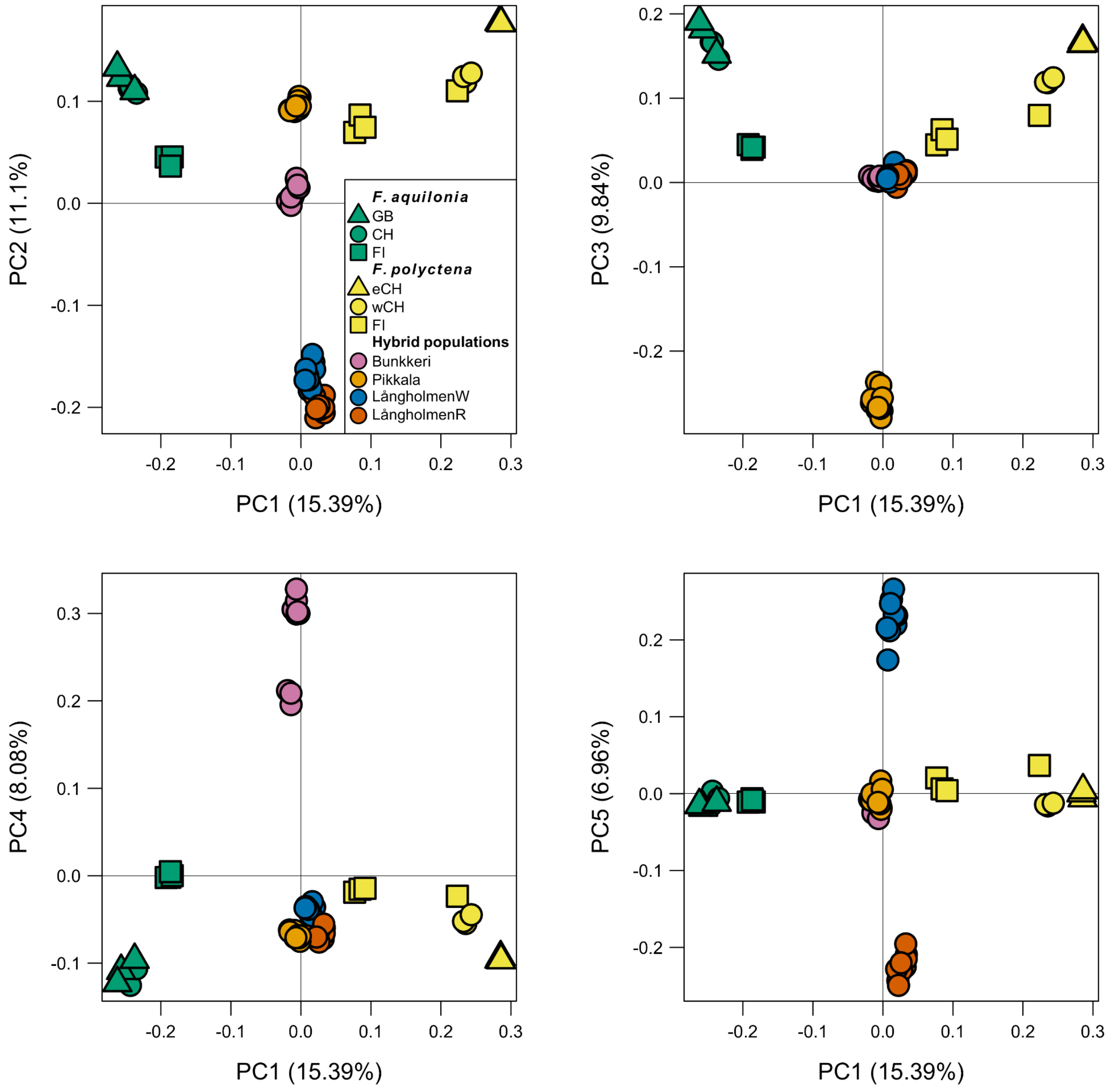


**Fig. S1.**

Visualization of the first five principal components of the PCA performed over 46,886 SNPs (5kb-thinned, Minor Allele Count ≥ 2).

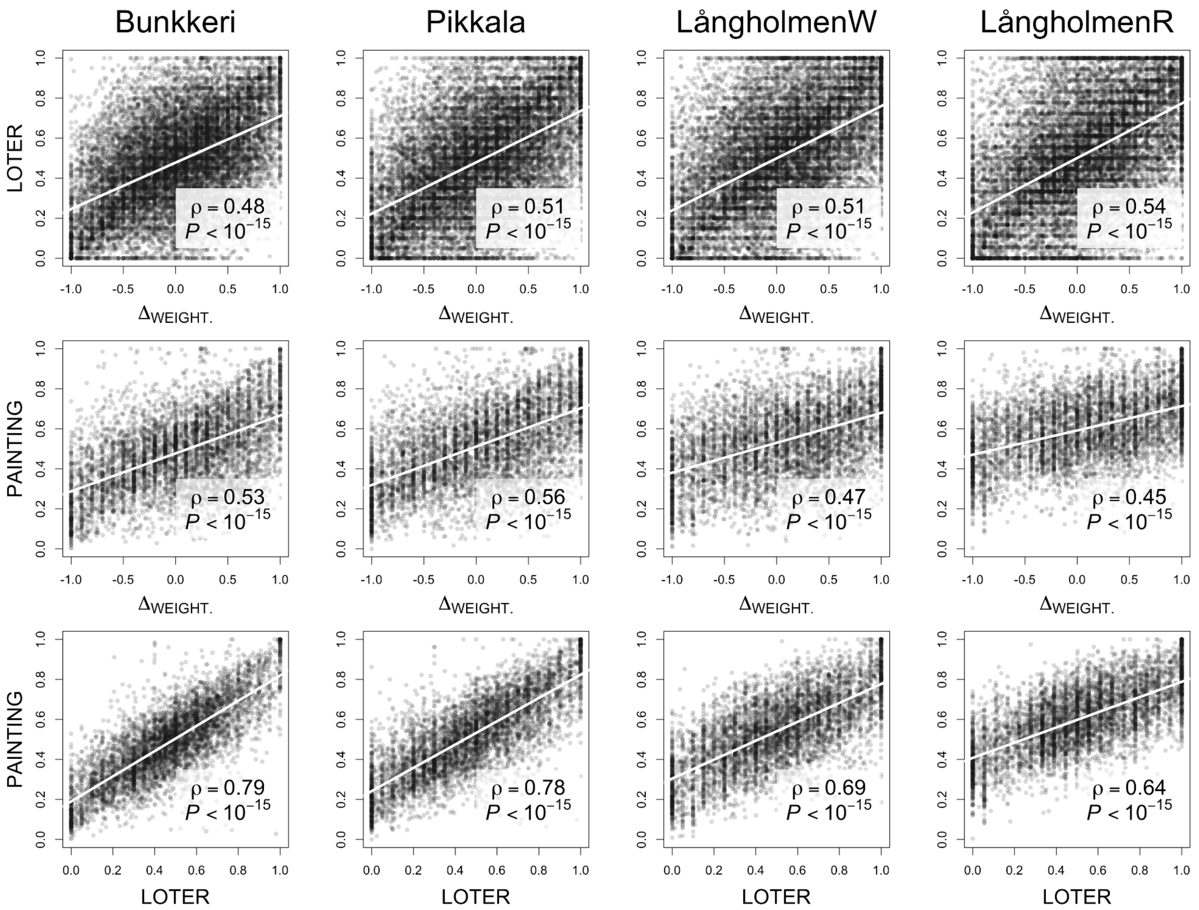


**Fig. S2.**

Comparison of ancestry mapping approaches. For each hybrid population (columns) are shown TWISST $\Delta_{\text{WEIGHT.}}$ statistics vs. LOTER local ancestry estimates (first row, 14,890 100-SNP windows), TWISST $\Delta_{\text{WEIGHT.}}$ statistics vs. naive chromosome painting local ancestry estimates (PAINTING, second row, 5,529 windows with at least five ancestry-informative SNPs), and LOTER vs. naive chromosome painting local ancestry estimates (third row, 5,529 windows with at least five ancestry-informative SNPs). $\Delta_{\text{WEIGHT.}}$ ranges between -1 if all topologies in the window group the hybrid population with *F. aquilonia*, to +1 if with *F. polyctena*. LOTER and naive chromosome painting are both SNP-based (results averaged over windows) and code ancestries as 0 for *F. aquilonia* and 1 for *F. polyctena*.
In each panel, the regression line is indicated in white. ρ, Spearman's correlation coefficient and *P*, *P*-value of the Spearman's correlation test.

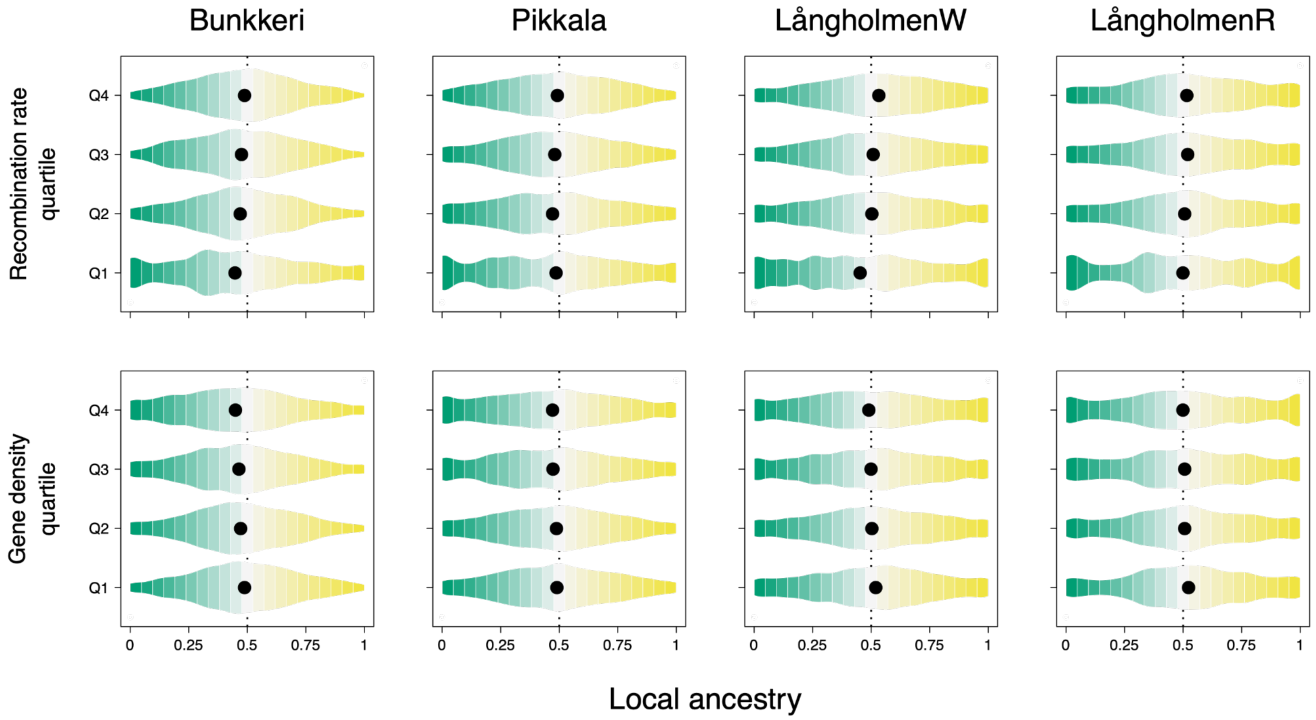


**Fig. S3.**

Distribution of LOTER local ancestry estimates (x-axis, 0: fixed for *F. aquilonia* ancestry component, 1: fixed for *F. polyctena* ancestry component) across recombination rate (upper row) and gene density (lower row) quartiles in each hybrid population (columns), computed over 20 kbp non-overlapping windows. Medians are indicated with black dots.

**Table S1.**

Sample information, sequencing statistics and accession numbers. In the “caste” column, w: worker and q: young unmated queen.

<see Excel file >

**Table S2.**

Demographic parameters estimated by fastsimcoal2 in demographic model analyses. Unless bounded, the upper limit of the search range could be exceeded. Each model used only a subset of these parameters. The time of admixture parameter (TADMS) indicated with an asterisk (*)  was first unconstrained, and then constrained to test for recent hybridization (< 50 generations). Double asterisks (**) mark parameters which calculation changes between models. The alternative minimum and maximum bounds are displayed in the respective columns.

<see Excel file >

**Table S3.**

Maximum likelihood parameter estimates for all models concerning the history of the *F. aquilonia* × *F. polyctena* hybrid population sampled in Pikkala (contained 348,228 sites). All effective sizes ($N_e$) are given in number of haploids. Times are given in number of generations. Migration rates are scaled according to population effective sizes (2Nm). Maximum-likelihood estimates for parameters are taken from the run reaching the highest composite likelihood of the 100 runs performed. Likelihoods are given in logarithmic scale. Maximum observed likelihood for this dataset is -1,377,703.973. ΔLikelihood is calculated by subtracting the expected likelihood from the maximum observed likelihood.

<see Excel file >

**Table S4.**

Maximum likelihood parameter estimates for all models concerning the history of the *F. aquilonia* × *F. polyctena* hybrid population sampled in Bunkkeri (contained 463,401 sites). All effective sizes ($N_e$) are given in number of haploids. Times are given in number of generations. Migration rates are scaled according to population effective sizes (2Nm). Maximum-likelihood estimates for parameters are taken from the run reaching the highest composite likelihood of the 100 runs performed. Likelihoods are given in logarithmic scale. Maximum observed likelihood for this dataset is -1,821,814.189. ΔLikelihood is calculated by subtracting the expected likelihood from the maximum observed likelihood.

<see Excel file >

**Table S5.**
Maximum likelihood parameter estimates for all models concerning the history of the *F. aquilonia* × *F. polyctena* hybrid population sampled in LångholmenW (contained 289,948 sites). All effective sizes ($N_e$) are given in number of haploids. Times are given in number of generations. Migration rates are scaled according to population effective sizes (2Nm). Maximum-likelihood estimates for parameters are taken from the run reaching the highest composite likelihood of the 100 runs performed. Likelihoods are given in logarithmic scale. Maximum observed likelihood for this dataset is -1,144,658.203. ΔLikelihood is calculated by subtracting the expected likelihood from the maximum observed likelihood.

<see Excel file >

**Table S6.**
Maximum likelihood parameter estimates for all models concerning the history of the *F. aquilonia* × *F. polyctena* hybrid population sampled in LångholmenR (contained 223,927 sites). All effective sizes ($N_e$) are given in number of haploids. Times are given in number of generations. Migration rates are scaled according to population effective sizes (2Nm). Maximum-likelihood estimates for parameters are taken from the run reaching the highest composite likelihood of the 100 runs performed. Likelihoods are given in logarithmic scale. Maximum observed likelihood for this dataset is -883,471.568. ΔLikelihood is calculated by subtracting the expected likelihood from the maximum observed likelihood.

<see Excel file >

**Table S7.**
Demographic parameters estimated by fastsimcoal2 in demographic model analyses. Unless bounded, the upper limit of the search range could be exceeded. Each model used only a subset of these parameters.

<see Excel file >

**Table S8.**
Maximum likelihood parameter estimates for the models concerning the history of the *F. aquilonia* × *F. polyctena* hybrid populations sampled in Bunkkeri and Pikkala (contained 328,913 sites). All effective sizes ($N_e$) are given in number of haploids. Times are given in number of generations. Migration rates are scaled according to population effective sizes (2Nm). Maximum-likelihood estimates for parameters are taken from the run reaching the highest composite likelihood of the 100 runs performed. Likelihoods are given in logarithmic scale. Maximum observed likelihood for this dataset is -1,509,459.108. ΔLikelihood is calculated by subtracting the expected likelihood from the maximum observed likelihood.

<see Excel file >

**Table S9.**
Maximum likelihood parameter estimates for all models concerning the history of the *F. aquilonia* × *F. polyctena* hybrid population sampled in Bunkkeri and LångholmenW (contained 282,215 sites). All effective sizes ($N_e$) are given in number of haploids. Times are given in number of generations. Migration rates are scaled according to population effective sizes (2Nm). Maximum-likelihood estimates for parameters are taken from the run reaching the highest composite likelihood of the 100 runs performed.
Likelihoods are given in logarithmic scale. Maximum observed likelihood for this dataset is -1,291,308.181. ΔLikelihood is calculated by subtracting the expected likelihood from the maximum observed likelihood.

<see Excel file >

**Table S10.**
Maximum likelihood parameter estimates for all models concerning the history of the *F. aquilonia* × *F. polyctena* hybrid population sampled in Pikkala and LångholmenW (contained 218,545 sites). All effective sizes ($N_e$) are given in number of haploids. Times are given in number of generations. Migration rates are scaled according to population effective sizes (2Nm). Maximum-likelihood estimates for parameters are taken from the run reaching the highest composite likelihood of the 100 runs performed.
Likelihoods are given in logarithmic scale. Maximum observed likelihood for this dataset is -1,001,623.462. ΔLikelihood is calculated by subtracting the expected likelihood from the maximum observed likelihood.

<see Excel file >

**Table S11.**
Maximum likelihood parameter estimates for all models concerning the history of the *F. aquilonia* × *F. polyctena* hybrid population sampled in LångholmenW and LångholmenR (contained 150,207 sites). All effective sizes ($N_e$) are given in number of haploids. Times are given in number of generations. Migration rates are scaled according to population effective sizes (2Nm). Maximum-likelihood estimates for parameters are taken from the run reaching the highest composite likelihood of the 100 runs performed.
Likelihoods are given in logarithmic scale. Maximum observed likelihood for this dataset is -693,262.061. ΔLikelihood is calculated by subtracting the expected likelihood from the maximum observed likelihood.

<see Excel file >
